# *Salmonella* Typhimurium infection drives NK cell loss and conversion to ILC1-like cells, and CIS inhibition enhances antibacterial immunity

**DOI:** 10.1101/2021.11.29.470332

**Authors:** Timothy R. McCulloch, Gustavo R. Rossi, Timothy J. Wells, Fernando Souza-Fonseca-Guimaraes

**Affiliations:** University of Queensland Diamantina Institute, The University of Queensland, Woolloongabba, QLD 4102, Australia; Australian Infectious Diseases Research Centre, University of Queensland, Brisbane 4072, QLD, Australia

**Keywords:** CIS, TGF-bR signaling, anti-bacterial responses, natural killer cells, innate lymphoid cell 1, cellular plasticity

## Abstract

Immunotherapy has revolutionized cancer therapy by reactivating tumor-resident cytotoxic lymphocytes. More recently, immunotherapy has emerged to restore immunity against infectious agents, including bacterial infections. Immunotherapy primarily targets inhibitory pathways in tumor-resident T cells, however interest in other effector populations, such as natural killer (NK) cells, is growing. We have previously discovered that NK cell metabolism, proliferation, and activation can be neutralized through the TGF-β immunosuppressive pathway by inducing plasticity of NK cells and differentiation into ILC1-like subsets. NK cells are also regulated through cytokine-inducible SH2-containing protein (CIS), which is induced by IL-15 and is a potent intracellular checkpoint suppressing NK cell survival and function. Targeting these two distinct pathways to restore NK cell function has shown promise is cancer models, but their application in bacterial infection remains unknown. Here, we investigate whether enhancement of NK cell function can improve anti-bacterial immunity, using *Salmonella* Typhimurium as a model. We identified conversion of NK cells to ILC1-like for the first time in the context of bacterial infection, however TGF-β signaling was curiously redundant in this plasticity. Future work should focus on identifying drivers of ILC1 plasticity and its functional implication in bacterial infection models. We further describe that CIS-deficient mice displayed enhanced pro-inflammatory function and dramatically enhanced anti-infection immunity. Inhibition of CIS may present as a viable therapeutic option to enhance immunity towards bacterial infection.

## Introduction

Natural killer (NK) cells are cytotoxic innate lymphocytes which have well described roles in the host defense against viral pathogens and cancer. Yet their ability to contribute to immunity in many bacterial infections, including *Salmonella enterica* serovar Typhimurium, remains unclear. NK cells have the potential to promote anti-*Salmonella* immunity through direct killing of infected cells and activation of infected cells through proinflammatory cytokine production, namely interferon (IFN)-γ. Previous studies in a murine model of *S.* Typhimurium infection have indicated that NK cells can contribute to protective IFN-γ and disease clearance in immunocompromised mice, but are otherwise dispensable when CD4^+^ T cells are present (Kupz et al., 2013). The idea of NK cell redundancy when adaptive lymphocytes are present has also been suggested in human disease (Vély et al., 2016). This suggests that the standard depletion model using anti-NK1.1 or anti-asialo-GM1 antibodies is insufficient to gauge the full potential of NK cells to promote antibacterial immunity in otherwise immunocompetent hosts. Depletion studies do not account for the impact of NK cell regulation during infection through the action of immunoregulatory cytokines such as interleukin (IL)-10 and transforming growth factor (TGF)-β, NK cell lymphopenia, and inhibitory immune checkpoints. Targeting these mechanisms of regulation may allow NK cells to participate in immunity where they may otherwise have been incapable.

Blockade of immunoregulatory molecules though immunotherapy have revolutionized cancer therapy. The classical targets are surface immune checkpoint molecules, such as programmed cell death protein-1 (PD-1), and cytotoxic T-lymphocyte-associated protein-4 (CTLA-4), which bind to their ligand on infected or tumor cells to ablate lymphocyte function. Blockade of these receptors can reverse this immune suppression and restore lymphocyte function (Robert, 2020). However, interest is rising in targeting other types of molecules, including regulatory cytokines and intracellular immune checkpoints. TGF-β is a pleiotropic cytokine of the TGF superfamily which has potent regulatory effects on NK cells by repressing mammalian target of rapamycin (mTOR) (Viel et al., 2016) and converting them to an innate lymphoid cell (ILC)1-like phenotype (Gao et al., 2017). Also of particular importance in NK cells is cytokine-inducible SH2-containing protein (CIS), which acts as a negative regulator of IL-15 signaling to limit NK cell proliferation and pro-inflammatory function (Delconte et al., 2016). Inhibition of these two pathways in bacterial infection may restore the function of NK cells, allowing them to contribute towards anti-bacterial immunity.

In the face of antimicrobial resistance, which is leading to infections that are increasingly difficult to treat, immunotherapy is arising as a potential alternative or conjunction to traditional antimicrobial therapy (McCulloch et al., 2021). In this study, we use a ‘gain of function’ model to investigate whether NK cell function could be enhanced or restored during infection to boost anti-bacterial immunity. Our group and others have focused on the improvement of NK cell function through the deletion of receptors for immunoregulatory molecules TGF-β (Gao et al., 2017; Rautela et al., 2019; Viel et al., 2016) or CIS (Delconte et al., 2016). Here, we investigate whether simultaneous immune checkpoint suppression of TGF-β and CIS signaling in a new transgenic mouse model could act synergistically to increase the magnitude of NK cell effector function against *Salmonella* infection.

## Results

### NK cells are depleted and converted to ILC1-like cells during *S.* Typhimurium infection

To investigate NK cells during *S*. Typhimurium infection, mice were infected i.p. with the attenuated mutant, SL32621. This model replicates the chronic, invasive infection seen in human disease. At day 10 post infection, spleens and livers were removed to observe phenotypic changes in immune cells. Of note, we observed a phenotypic switch from conventional NK (cNK) cells to ILC1-like cells, as determined by upregulation of tissue residence marker CD49a. This occurred in both the spleen (**Fig. 1a,b**) and liver (**Fig. 1c,d**) of infected mice. Furthermore, we also observed significant NK cell lymphopenia in the blood (**Fig. 1e**) and spleen (**Fig. 1f**) of infected mice compared to uninfected. NK cell lymphopenia was not seen in the liver of infected mice, where normal NK numbers were persevered (**Fig. 1g**). Collectively, this data indicates that NK cells are considerably affected during *S.* Typhimurium infection, characterized by conversion to ILC1-like cells and organ specific depletion.

**Figure 1:**
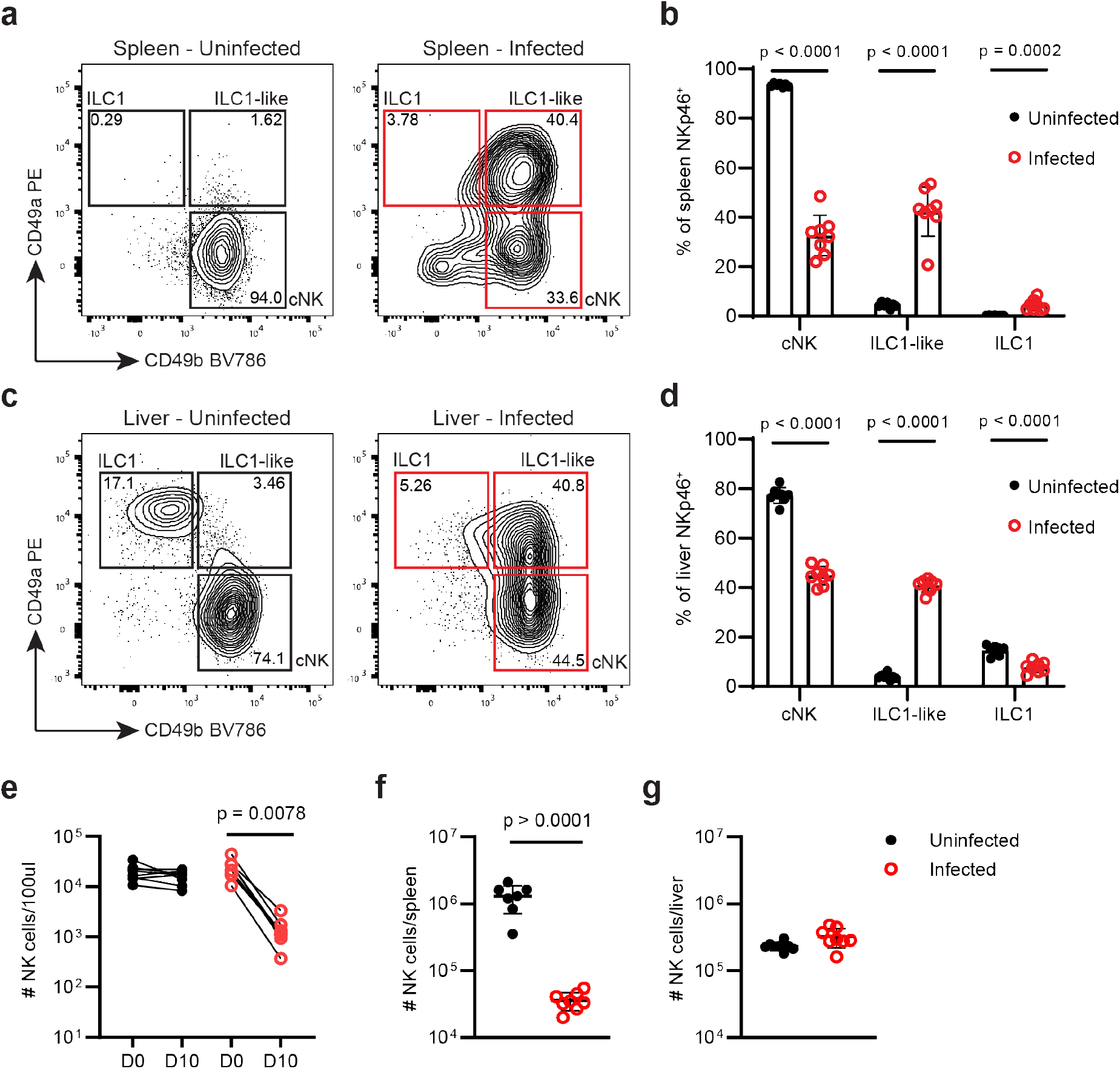
NK cells are depleted and converted to ILC1-like cells during *S.* Typhimurium infection. Group 1 ILCs were analysed by flow cytometry 10 days after infection with *S.* Typhimurium. Representative flow cytometry plots showing CD49a and CD49b expression from one uninfected and one infected mouse spleen (**a**) and liver (**c**) are shown, along with graphs displaying relative percentages of cNK, ILC1-like, and ILC1 within the spleen (**b**) and liver (**d**) (n = 8). Total numbers of NK cells per 100μl of blood prior to infection and at day 10 post infection were determined by flow cytometry (**e**). At 10 days post infection, numbers of NK cells in the spleen (**f**) and liver (**g**) were determined by flow cytometry. Data from one experiment. Each symbol represents an individual mouse, graphs show mean value ± SEM. Statistical p values determined by Mann-Whitney *t* test, or Wilcoxon rank test for (**e**). No p value indicates no significant difference.

### Deletion of *CISH*, but not conditional deletion of *TgfbRII*, results in enhanced anti-bacterial immunity

Conversion of NK cells to ILC1-like cells has previously been shown to contribute to immune evasion of tumors (Gao et al., 2017), where the conversion is driven primarily by TGF-β (Gao et al., 2017; Viel et al., 2016). Thus, we predicted that conversion to ILC1-like cells in our infection model could be dampening the ability of NK cells to contribute to bacterial clearance. To address this, we infected mice with conditional deletion of the TGF-β receptor II specifically within NK cells (*TgfbRII*^*FL*^). NK cell mediated bacterial clearance could be further impacted by the observed lymphopenia. Deletion of the *Cish* gene, which encodes for the intracellular checkpoint molecule CIS, has been reported to enhance NK cell function and proliferation (Delconte et al., 2016, 2020). While deletion of CIS does not lead to increased NK cell accumulation in the steady state (Delconte et al., 2016), we predicted its deletion could enhance proliferation to help maintain NK cell numbers during infection. Therefore, we also infected CIS deficient mice (*Cish*^*KO*^). We also infected mice with CIS deficiency and conditional deletion of TGF-β signaling within NK cells (*TgfbRII*^*FL*^/*Cish*^*KO*^) to determine if these could have a synergistic effect in improving anti-bacterial immunity. Surprisingly, *TgfbRII*^*FL*^ mice did not show a reduced bacterial load in either the spleen (**Fig. 2a**) or the liver (**Fig. 2b**) compared to wild-type controls. Conversely, *Cish*^*KO*^ mice exhibited a significant reduction in bacterial burden in both organs (**Fig, 2a,b**). The combination of both NK cell gain of function genes did not synergize further reduce bacterial burdens (**Fig, 2a,b**). To investigate whether the enhanced immunity in *Cish*^*KO*^ mice was NK cell dependent, *Cish*^*KO*^ mice were treated with αNK1.1 to deplete NK cells. No differences were observed in the spleen (**Figure, 2c**). In the liver, while *Cish*^*KO*^ was able to significantly reduce bacterial burdens, *Cish*^*KO*^ with NK cell depletion did not significantly differ from wild-type or *Cish*^*KO*^ alone (**Figure, 2d**). Further, IL-6 was the only cytokine significantly increased in the plasma of *Cish*^*KO*^ mice at day two compared to wild-type controls (**Supplementary Figure 1**), suggesting *Cish* deletion may primarily act through myeloid cells in this case. No significant increases in cytokine levels were observed at day nine post infection (**Supplementary Figure 2**). Thus, we found no evidence that the reductions in bacterial burdens observed in the livers of *Cish*^*KO*^ mice was due to enhancement of NK cell function.

**Figure 2:**
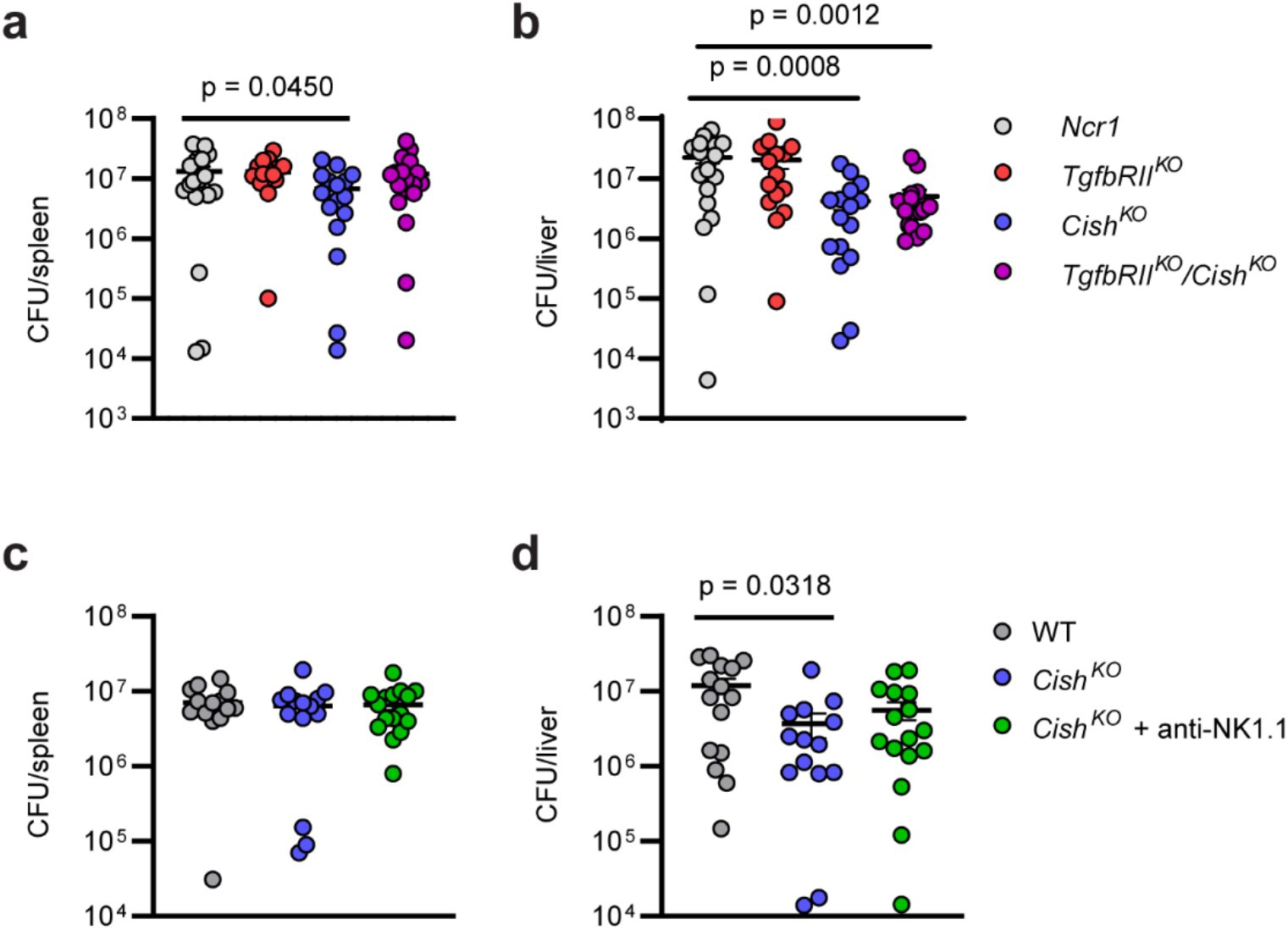
Deletion of *CISH*, but not conditional deletion of TGF-β signalling, results in reduced bacterial burdens. *Ncr1Cre, TgfbRII*^*FL*^, *Cish*^*KO*^, and *TgfbRII*^*FL*^/*Cish*^*KO*^ mice were infected with *S.* Typhimurium. At day 10 post infection, spleens and livers were collected to quantify bacterial burden. Colony forming units (CFU) per spleen (**a**) and liver (**b**) are shown (n = 15-19). Data from two independent experiments. C57BL/6 and *Cish*^*KO*^ were infected while simultaneously treated with either anti-NK1.1 depletion antibody, or isotype control. Colony forming units per spleen (**c**) and liver (**d**) are shown (n = 14-16). Data from two independent experiments. Each symbol represents an individual mouse, graphs show mean value ± SEM. Statistical p values determined by Mann-Whitney *t* test, where no p value indicates no significant difference.

### Conversion of NK to ILC1-like cells is not TGF-β dependent in *S.* Typhimurium infection

We found that NK cells are converted to ILC1-like cells during *S.* Typhimurium infection (**Fig 1a-d**), a plasticity which has been previously shown to be dependent primarily on TGF-β (Gao et al., 2017). However, despite this conversion being well characterized in the literature to contribute to immune evasion during cancer (Gao et al., 2017; Hawke et al., 2020; Viel et al., 2016), conditional deletion of TGF-β within NK cells did not appear to improve anti-bacterial immunity. We postulated this could be due to one of two reasons, either ILC1-like conversion does not restrict NK cell mediated immunity in our model, or TGF-β is not the sole driver of this conversion. To investigate this, the NK/ILC1-like status was investigated in *TgfbRII*^*FL*^ mice infected with *S.* Typhimurium by flow cytometry. Surprisingly, conditional deletion of TGF-β signaling within NK cells did not prevent conversion to ILC1-like cells in either the spleen (**Fig. 3a,b**) or liver (**Fig. 3c,d**) of infected mice. We also targeted TGF-β signaling therapeutically using the TGF-β receptor 1 kinase inhibitor galunisertib in infected mice, however we observed no changes to bacterial burdens (**Supplemental Fig. 3a,b**), weight change (**Supplemental Fig. 3c**), or NK cell to ILC1-like conversion (**Supplemental Fig. 3d,e**) in galunisertib treated mice compared to untreated controls. Further, neither conditional deletion of *TgfbrII*, deletion of *Cish*, or a combination were able to prevent NK cell lymphopenia observed in the blood and spleen of infected mice (data not shown). Taken together, this data suggests that canonical TGF-β / TGF-bRII signaling is redundant in driving NK cell to ILC1-like conversion during *S*. Typhimurium infection. This may explain why no reduction in bacterial burdens were observed in *TgfbRII*^*FL*^ mice or mice treated with the TGF-bRI inhibitor galunisertib, although whether the conversion limits NK-mediated immunity during bacterial infection remains unclear.

**Figure 3:**
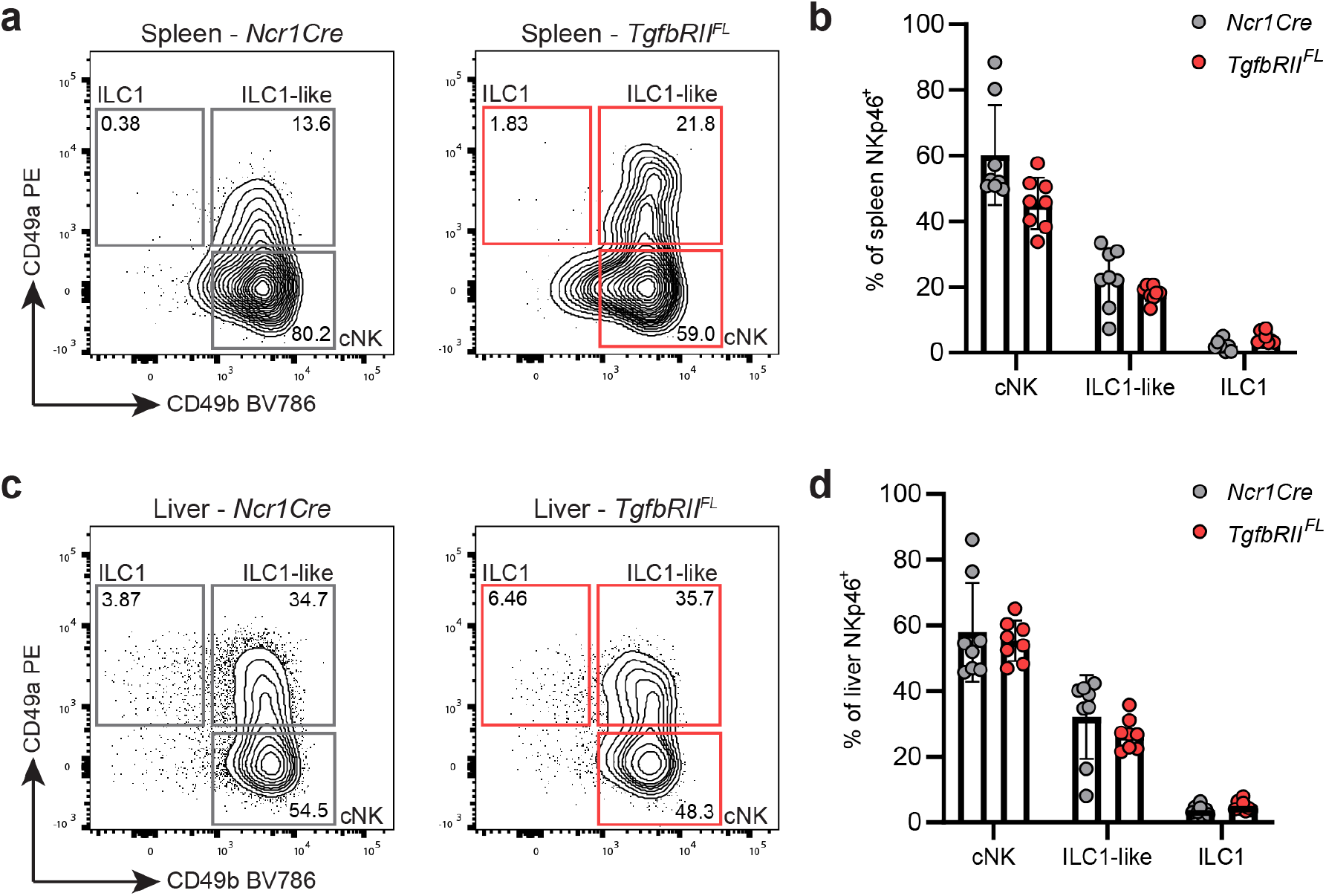
TGF-β signalling is redundant in NK to ILC1-like plasticity in *S.* Tyhpimurium infection. *TgfbRII*^*FL*^ mice, which lack the TGF-β receptor specifically within NK cells, as well as *Ncr1Cre* controls, were infected with *S.* Typhimurium. At day 10 post infection, Group 1 ILCs were analysed by flow cytometry. Representative flow cytometry plots showing CD49a and CD49b expression from one *Ncr1Cre* and one *TgfbRII*^*FL*^ mouse spleen (a) and liver (c) are shown, along with graphs displaying relative percentages of cNK, ILC1-like, and ILC1 within the spleen (b) and liver (d) (n = 8). Data from one experiment. Each symbol represents an individual mouse, graphs show mean value ± SEM. Statistical p values determined by Mann-Whitney *t* test, where no p value indicates no significant difference.

## Discussion

The purpose of this study was to gain a better understanding of how NK cells are regulated during bacterial infection. NK cells are increasingly being recognized as a promising immunotherapy target in settings of cancer (Souza-Fonseca-Guimaraes et al., 2019), however their efficacy as a target during acute or chronic bacterial infection is unknown. Immunotherapy for treating bacterial infections, particularly in the case of antibiotic resistance, is an emerging field (McCulloch et al., 2021), for which NK-mediated immunotherapy may also show potential. Identification of specific regulatory molecules and checkpoints acting on NK cells during bacterial infection could uncover novel immunotherapy targets to enhance NK-mediated bacterial immunity.

Our finding that NK cells were converted to ILC1-like cells during *S*. Typhimurium was not unexpected. This conversion has previously been observed in other diseases including cancer (Gao et al., 2017) and parasitic infection (Park et al., 2019). Further, *Salmonella* subspecies are known to actively drive macrophage polarization towards an anti-inflammatory (or M2-like) phenotype (Stapels et al., 2018), promoting the production of regulatory cytokines such as IL-10 and TGF-β (Jaslow et al., 2018). However, by using transgenic mice and therapeutic inhibition, we found that conversion to ILC1-like cells was likely not TGF-β dependent in our model. The TGF-β superfamily is a large group of over 33 regulatory proteins which have both distinct and overlapping functions (Morikawa et al., 2016). Therefore it is reasonable to assume that other members of the TGF-β superfamily could also drive this conversion, which has already been observed in the case of Activin A (Rautela et al., 2019). Our findings using the therapeutic inhibitor galunisertib, which targets the TGF-β receptor 1 kinase, may also be hindered by inconsistencies in the literature on the appropriate dosing of this drug in mouse models. Dosing regimens published in murine studies range from 10 mg/kg every second day by i.p. injection (Rautela et al., 2019) to 150 mg/kg twice daily by oral gavage (Gunderson et al., 2020). Whether underdosing was a defining factor in our results is unknown. The driving factor, or factors, behind NK to ILC1-like conversion would need to be identified before definitively determining whether this conversion limits anti-bacterial immunity.

This study also found that deletion of CIS improved control of *S*. Typhimurium infection. CIS is a suppressor of cytokine signaling (SOCS) protein which functions as a negative regulator of cytokine signaling. In the case of NK cells, IL-15 signaling leads to phosphorylation of Janus kinase (JAK)1 and 3, and subsequent recruitment and phosphorylation of STAT5. This allows STAT5 to then translocate to the nucleus to transcribe variety of genes associated with NK cell function and survival (Ma et al., 2006). Also transcribed is *Cish*, encoding CIS, which binds to JAK1 and JAK3, targeting them for proteasomal degradation. In this way, CIS acts as a negative feedback loop to limit IL-15 signaling and NK cell activation (Delconte et al., 2016). However, STAT5 signaling is also active in other cell types in response to signaling by cytokines other than IL-15, including IL-2 and GM-CSF. CIS also inhibits T cell receptor signaling (Palmer et al., 2015). Thus, CIS has also been shown to regulate additional immune cells including ILC2s (Kotas et al., 2021), CD4^+^ and CD8^+^ T cells (Palmer et al., 2015; Yang et al., 2013), neutrophils (Louis et al., 2020), and macrophages (Carow et al., 2017; E. et al., 2021). Considering this was a whole mouse knockout of CIS, we cannot be sure of which cell type, or combinations of cell types, benefited from CIS deletion to promote anti-bacterial immunity. We found no evidence to confirm our initial hypothesis that CIS deletion acted through NK cell enhancement.

Curiously, our results may be somewhat conflicting with other studies. Despite being a negative regulator of immune function, CIS has previously been shown to mediate early control of tuberculosis infection in a mouse model (Carow et al., 2017). In humans, single nucleotide polymorphisms (SNPs) in *Cish* have been associated with increased susceptibility to tuberculosis, malaria, and bacteremia (Khor et al., 2010). However, the functional implications of these SNPs on immune response were not established in the study. These previous results may be explained by multiple observations in which excessive inflammation worsens infectious disease. This can be epitomized by the curious case of anti-PD1 therapy in tuberculosis, which exacerbates disease severity and reactivates latent infection (Barber et al., 2019; Kauffman et al., 2021). Thus, in the case of tuberculosis, immune enhancement does not always lead to greater immunity. That we have found CIS deletion enhances anti-*Salmonella* immunity, where others have found it impedes anti-tuberculosis immunity, may be explained by underlying differences in the physiology and pathology of these two pathogens. These differences must be rigorously examined if any pharmacological targeting of CIS signaling is to be trialed to treat bacterial infections.

In summary, we have been able to expand the current knowledge of ILC plasticity to show for the first-time evidence of conversion of NK cells to ILC1-like cells during bacterial infection. However, the precise driver of conversion as well as the functional relevance this plasticity has on infection outcome remains elusive. Here, we have also shown that CIS is a potent immune checkpoint in anti-*Salmonella* immunity. However, expression of CIS is also conserved across other immune cell types such as myeloid cells, and thus the exact cell type or mechanism where CIS acts on to restrict bacterial clearance could not be addressed in this current study. Future work is warranted to elucidate how CIS inhibition enacts its effects, and whether pharmacological inhibition of this molecule could enhance anti-bacterial immunity.

## Methods

### Mouse models

*Ncr1*^*cre/wt*^*TgfbRII*^*fl/fl*^ mice were used as conditional TgfbRII-deficiency specific to NK cells, obtained by crossing as previous described by our group (Gao et al., 2017). *Cish*^*-/-*^ were maintained on a C57BL/6 background (Delconte et al., 2016). To obtain a double deficient mouse model, *Ncr1*^*cre/wt*^*TgfbRII*^*fl/fl*^ were back crossed with *Cish*^*-/-*^ to obtain a *Cish*^*-/-*^ *Ncr1*^*cre/wt*^*TgfbRII*^*fl/fl*^ mice strain. *Ncr1*^*cre/wt*^ mice were considered wild-type controls for some experiments. *Ncr1*^*cre/cre*^*Mcl1*^*fl/fl*^ mice were used as NK cell deficient controls (Sathe et al., 2014). All experiments were performed using cells from age/sex matched cohort of mice (age range 8-12 weeks). Cohort sizes are described in each figure legends, where power calculations were used to estimated sample size needed to achieve statistical significance at a 50% change in bacterial burden or immune parameters with a power of 0.80 and Type I error (alpha) of 0.05. For infection studies, mice with no detectable bacterial load at the end of the experiment were considered to have not taken up infection and were excluded from further analysis. All experiments were approved by the University of Queensland’s Animal Ethics Committees.

### Bacterial strains and *in vivo* infections

Mice were infected with an attenuated *aroA* mutant strain of *Salmonella enterica* serovar Typhimurium, SL3261 (Hoiseth & Stocker, 1981). For *in vivo* infection, bacteria were grown at 37°C with shaking in Lysogeny broth (LB) for 16 to 18 hours. OD_600_ was used to enumerate bacteria, before being diluted to the appropriate concentration in PBS. Mice were infected by intraperitoneal (i.p) injection with 5 × 10^6^ colony forming units (CFU) of SL3261 in 200μl and sacrificed at the described times post-infection. The TGF-β receptor I inhibitor galunisertib (LY2157299, SelleckChem, Houston TX) was given at a dose of 10mg/kg by i.p infection, as described previously (Rautela et al., 2019). To deplete NK cells, appropriate mice were treated with 100μg of anti-NK1.1 antibody (PK136, BioXCell, Lebanon NH) or isotype control (2AE, BioXCell) on days −3, 0, 3, and 8 relative to infection.

### Murine tissue collection

Blood samples were taken from mice by retro-orbital bleeds into EDTA-coated tubes. Tubes were centrifuged at 1,500 g for 15 minutes, and plasma removed from cell pellet. Plasma samples were stored at −20°C until analysis. At the experimental endpoint, mice were euthanized by CO_2_ asphyxiation. Spleens and livers were dissected and held in PBS until processing. Bacterial counts were enumerated from organs by homogenizing samples in 0.1% Triton-X (Sigma-Aldrich, Burlington MA) in PBS before serially diluting in PBS and plating on LB agar plates.

### Flow cytometry

Spleens and livers were passed through a 70μm or 100μm cell strainer, respectively, in cold fluorescence-activated cell sorting (FACS) buffer (PBS containing 2% fetal bovine serum and 2mM EDTA). Leukocytes were enriched using 37.5% Percoll solution (GE Healthcare, Uppsala, Sweden)) and red blood cells lysed with Ammonium-Chloride-Potassium (ACK) lysis buffer (Biolgend, San Diego CA). Dead cells were stained with Fixable Viability Stain 440UV (1:1000 in PBS, Becton Dickinson, Franklin Lakes NJ) for 15 minutes at room temperature. Fc receptors were blocked by incubation for 15 minutes in Fc Blocking Reagent (1:100 in FACS buffer, Miltenyi Biotec, Bergisch Gladbach, Germany). Single-cell suspensions were stained with the indicated fluorescent antibodies on ice for 45 minutes. For intracellular cytokine staining, cells were fixed and permeabilized using the FoxP3/Transcription Factor Staining Buffer Set (eBioscience, San Diego CA), then stained for 60 minutes with the indicated fluorescent antibodies. Antibodies targeting CD45 (30-F11), CD4 (GK1.5), CD3 (145-2C11), CD8a (53-6.7), TIGIT (1G9), CD69 (H1.2F3), CD314 (CX5), CD223 (C9B7W), CD49b (HMa2), CD11b (M1/70), CD44 (IM7), PD-1 (J43), CD49a (HA31/8) were purchased from Becton Dickinson (Franklin Lakes NK). Antibodies targeting CD226 (TX42.1), CD335 (29A1.4), KLRG1 (2F1/KLRG1), CD19 (6D5), Ly6G (1A8), F4/80 (BM8), and CD62L (MEL-14) were purchased from Biolegend. Antibodies targeting Tim-3 (RMT3-23), Eomes (Dan11mag), and FoxP3 (FJK-16S) were purchased from eBioscience (San Diego CA).

### Measurement of cytokines

IFN-γ titers were determined from murine serum samples using a Mouse IFN-γ ELISA set (Becton Dickinson) as per the manufacturer’s instructions. Other cytokines were determined using Cytometric Bead Array (Becton Dickinson) as per the manufacturer’s instructions.

### Statistics

Statistical analysis was performed using GraphPad Prism Software v9 (GraphPad Software, San Diego CA). Statistical tests were performed for experiments as indicated in figure legends, and error bars represent SEM. Levels of statistical significance are expressed as p values.

## Author Contributions

T.R.M. and F.S.F.G. designed research and wrote the paper. T.R.M., and G.R.R., performed research and analysed data. T.J.W and F.S.F.G. supervised work.

## Acknowledgments

We thank all the members of the Guimaraes and Wells laboratories; Prof. N. D. Huntington, Prof. G. T. Belz, Prof. S. Bell, Prof. M. Sweet, and Prof. A. Yoshimura for discussion, comments, and advice in this project; Prof. E. Vivier for providing the NKp46cre mice; and Profs. J. Ihle and E. Parganas for providing the CIS knockout mice. This research was carried out at the Translational Research Institute, Woolloongabba, QLD 4102, Australia. The Translational Research Institute is supported by grants from the Australian and Queensland Governments.

## Supplementary Material

**Figure 2 – figure supplement 1:**
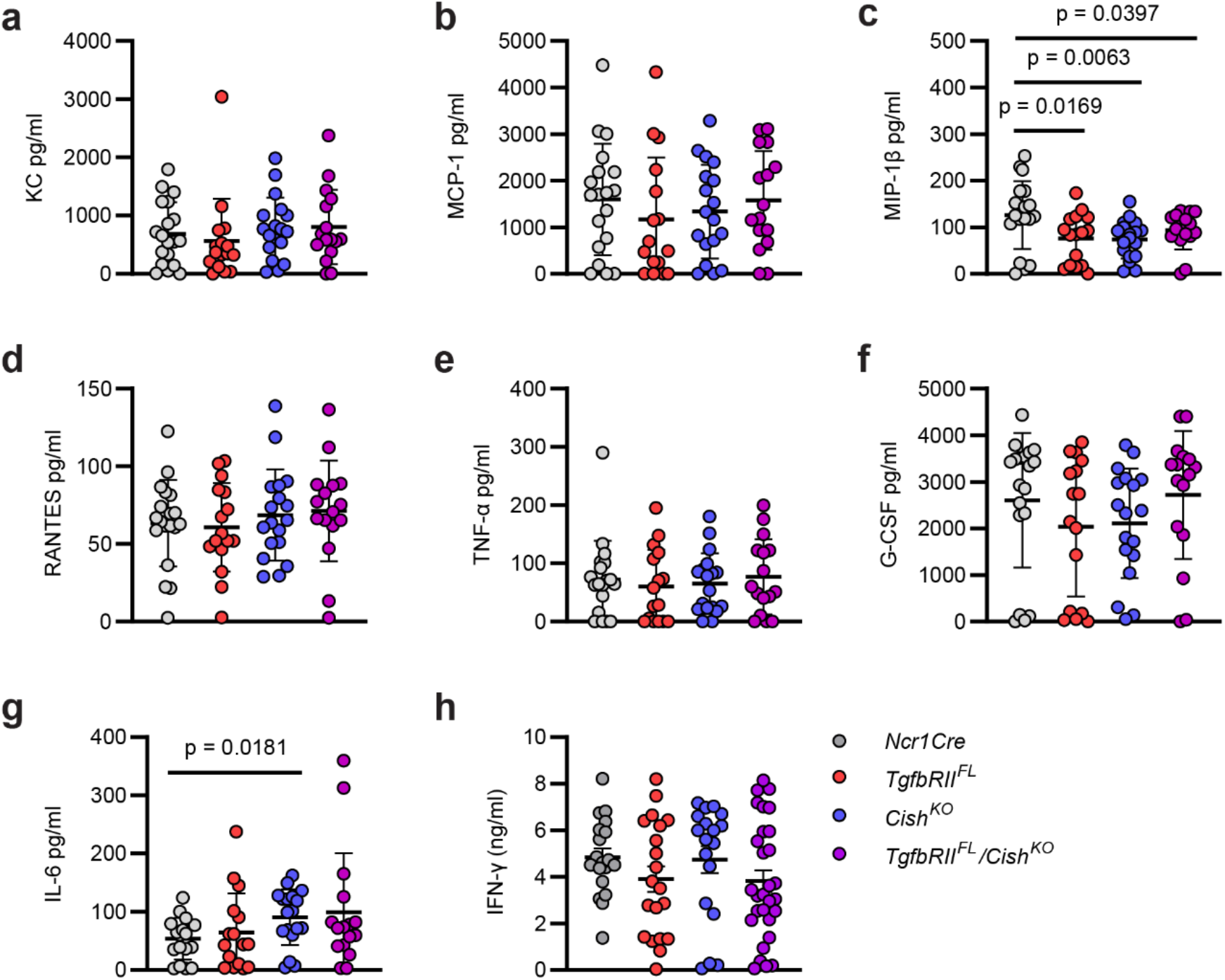
Plasma cytokine levels in transgenic mice at day 2 post infection. *Ncr1Cre, TgfbRII*^*FL*^, *Cish*^*KO*^, and *TgfbRII*^*FL*^/*Cish*^*KO*^ mice were infected with *S.* Typhimurium. At day 2, plasma was taken to measure cytokine levels by CBA (**a-g**) or ELISA (**h**). Levels of KC (**a**), MCP-1 (**b**), MIP-1β (**c**), RANTES (**d**), TNF-α (**e**), G-CSF (**f**), IL-6 (**g**), and IFN-γ (**h**) are shown. Data from two individual experiments. Each symbol represents an individual mouse, graphs show mean value ± SEM. Statistical p values determined by Mann-Whitney *t* test, where no p value indicates no significant difference.

**Figure 2 – figure supplement 2:**
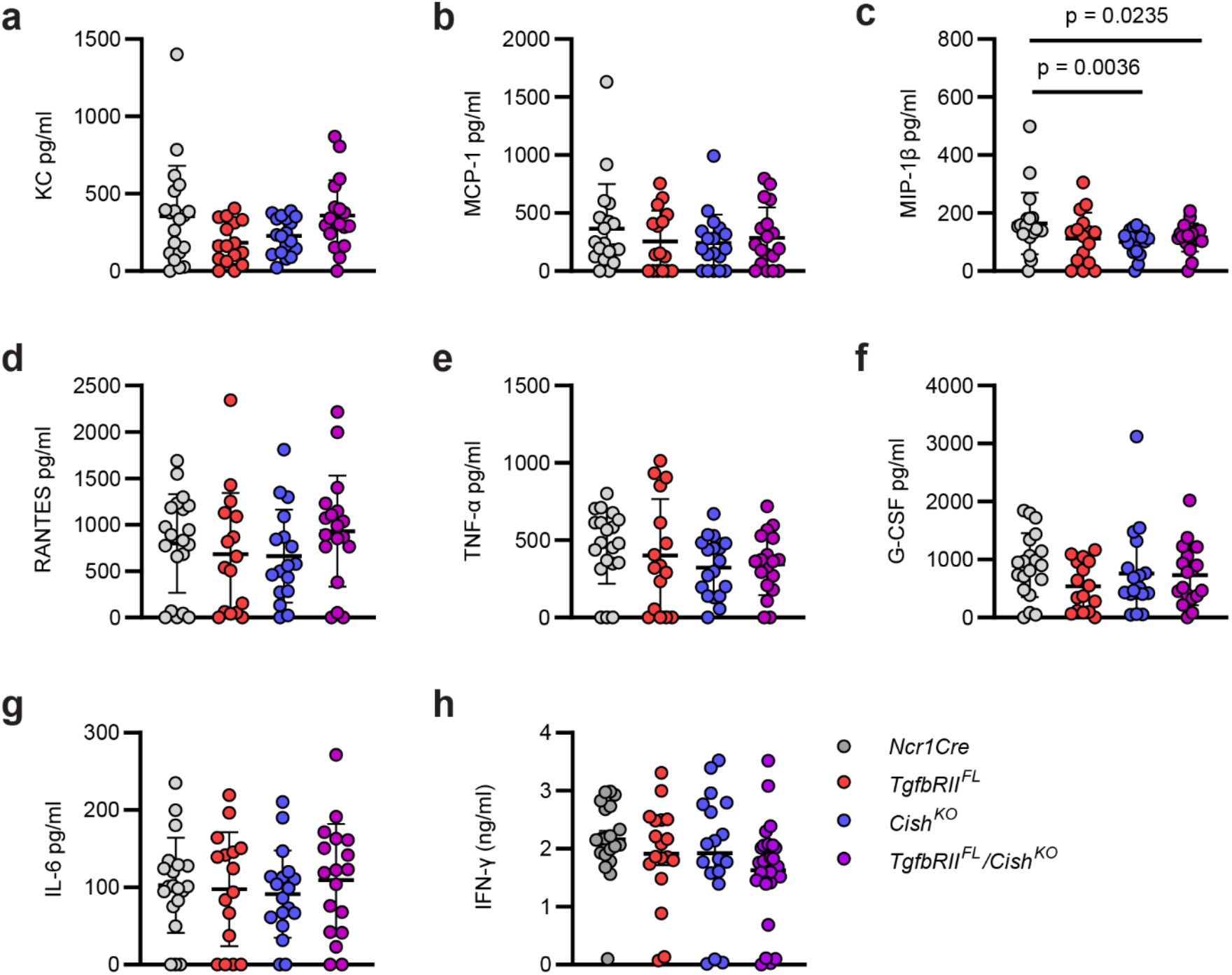
Plasma cytokine levels in transgenic mice at day 9 post infection. *Ncr1Cre, TgfbRII*^*FL*^, *Cish*^*KO*^, and *TgfbRII*^*FL*^/*Cish*^*KO*^ mice were infected with *S.* Typhimurium. At day 9, plasma was taken to measure cytokine levels by CBA (**a-g**) or ELISA (**h**). Levels of KC (**a**), MCP-1 (**b**), MIP-1β (**c**), RANTES (**d**), TNF-α (**e**), G-CSF (**f**), IL-6 (**g**), and IFN-γ (**h**) are shown. Data from two individual experiments. Each symbol represents an individual mouse, graphs show mean value ± SEM. Statistical p values determined by Mann-Whitney *t* test, where no p value indicates no significant difference.

**Figure 3 – figure supplement 1:**
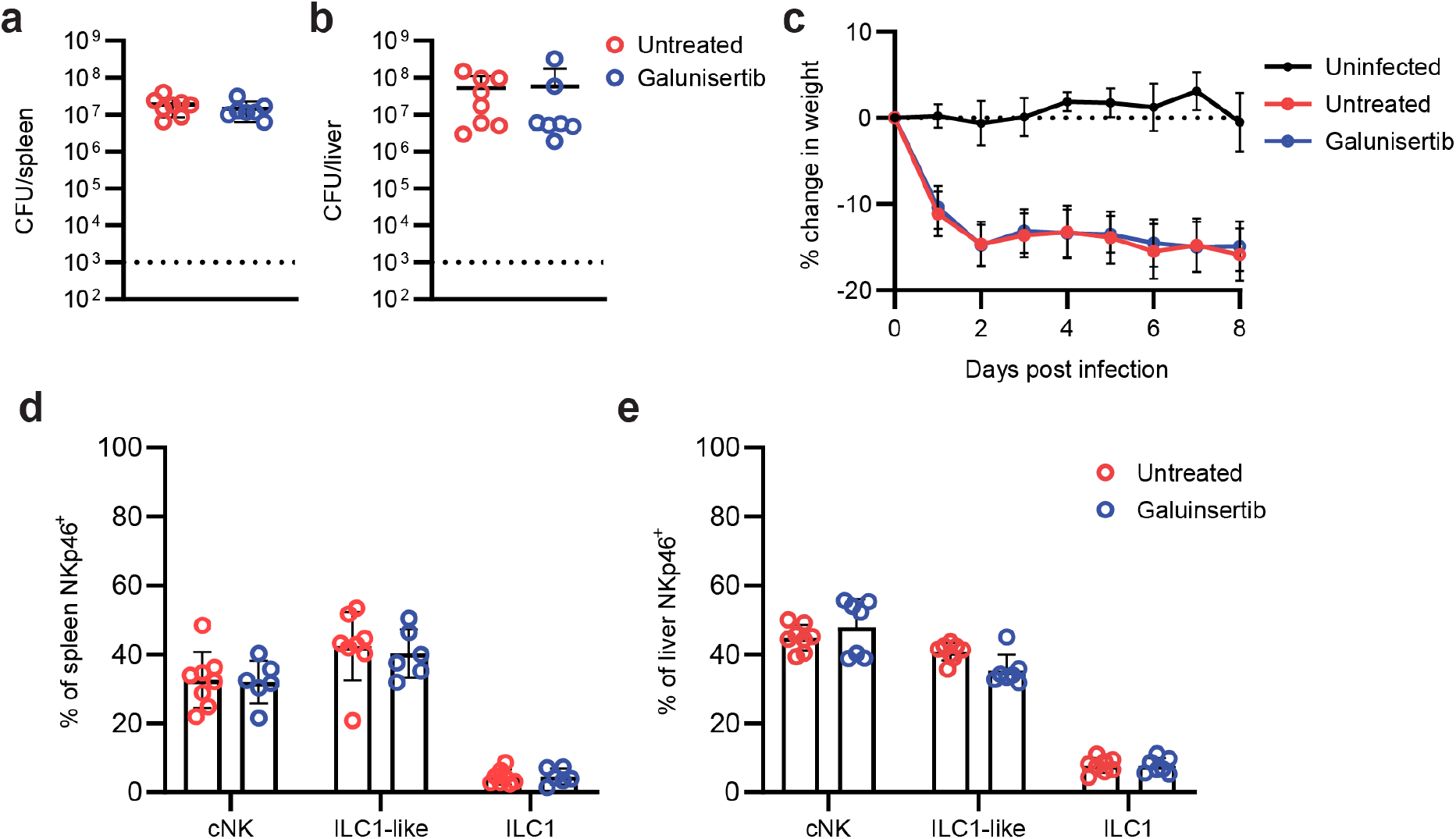
Galunisertib has no effect on *S*. Typhimurium infection. C57BL/6 mice were infected with *S*. Typhimurium and treated every second day with the TGF-β antagonist, galunisertib. At day 10, spleens and livers were removed to quantify bacterial burdens and immune parameters. CFU per spleen (**a**) and liver (**b**) are shown, along with % change in weight over the course of infection (**c**). Flow cytometry was performed to determine relative percentages of cNK, ILC1-like, or ILC1 within NKp46^+^ cells in the spleen (**d**) and liver (**e**)(n = 7-8). Data from one experiment. Each symbol represents an individual mouse, graphs show mean value ± SEM. Statistical p values determined by Mann-Whitney *t* test, where no p value indicates no significant difference.

